# Copy number of the *bla*_NDM-1_ gene (New Delhi metallo-β-lactamase-1) and meropenem susceptibility in Enterobacterales: Consideration for treatment with meropenem?

**DOI:** 10.1101/2025.09.03.673909

**Authors:** Amrita Bhattacharjee, Soumalya Banerjee, Sanjib Das, Tanusree Ray, Nibendu Mondal, Agniva Majumdar, Sulagna Basu

**Affiliations:** Division of Bacteriology, ICMR-National Institute for Research in Bacterial Infections (Formerly ICMR-NICED), P-33 CIT Road, Scheme XM, Beliaghta, Kolkata-700010, West Bengal, India; Department of Biological Sciences, Bose Institute, Kolkata-700091, West Bengal, India; Department of Microbiology & Cell Science, Fort Lauderdale Research and Education Center, University of Florida, Davie, Florida 33314, USA; Regional Virus Research and Diagnostic Laboratory (VRDL), ICMR-National Institute for Research in Bacterial Infections (Formerly ICMR-NICED), P-33 CIT Road, Scheme XM, Beliaghta, Kolkata-700010, West Bengal, India

**Keywords:** *bla*_NDM_ expression, ddPCR, meropenem efficacy, neutropenic mice, time-kill assay

## Abstract

Some Enterobacterales carrying *bla*_NDM_ were still susceptible to meropenem, exhibiting meropenem minimum inhibitory concentration (MIC) values below the clinical breakpoint. As this phenomenon is a rare observation, this study investigated the underlying regulatory mechanisms contributing to the low MIC values and evaluated the efficacy of meropenem as a therapeutic preference for such isolates. Six clinical *bla*_NDM_-positive Enterobacterales exhibited meropenem MICs between 0.5-2.0 mg/L. Successful transmission of *bla*_NDM_ into recipient *E. coli* J53 indicated its presence on plasmids. Conventional Sanger and whole-genome sequencing revealed that no truncation or deletion was observed within *bla*_NDM_ or its core promoter region up to 120 bp upstream. Quantitative real-time PCR of *bla*_NDM_ showed significantly lower expression levels of *bla*_NDM_, i.e., 0.2-0.5 in study isolates, compared to a clinical *bla*_NDM_-possessing isolate (expression=1) exhibiting a high meropenem MIC value (64 mg/L). Droplet digital PCR (ddPCR) further demonstrated that these isolates carried a lower copy number of the *bla*_NDM_ gene, compared to the *bla*_NDM_-possessing isolate with high meropenem MIC (64 mg/L). Furthermore, the *in vitro* time-kill assay exhibited bactericidal efficacy of meropenem >3 log10 decrease in cfu/mL over the 24 hours from the starting inoculum. Administration of meropenem in *in vivo* neutropenic mice produced a 4-log cfu reduction compared to untreated controls at the start of dosing. Our data suggest that a reduction in *bla*_NDM_ gene dosage, reflecting lower expression of *bla*_NDM_, correlates with lower meropenem MIC values. Meropenem is effective against these isolates when the meropenem MICs for the isolates are 0.5-2.0 mg/L.

**Importance:** The production of NDM by Enterobacterales poses one of the most urgent global public health threats due to its plasmid-mediated spread, leading to treatment failure by exhibiting high MICs against a broad-spectrum of antibiotics. However, in rare instances, *bla*_NDM-1_-positive clinical Enterobacterales with unexpectedly low meropenem MICs (0.5–2.0 mg/L) raise a further question of reconsidering meropenem as a potential therapeutic option in these specific cases. This study identified that low MICs were associated with reduced expression and lower copy number of the plasmid-mediated *bla*_NDM_ gene, despite intact promoter regions. Both *in vitro* and *in vivo* assays validated the bactericidal activity of meropenem against these isolates. The study raised an argument about the potential limitations of relying solely on genotypic detection of resistance genes for therapeutic decision-making, which emphasises the need for further investigation. As a safe medication, repurposing meropenem for neonatal treatment may eliminate the need for other toxic drugs.

## Introduction

Carbapenems are the broadest spectrum of antibacterial agents in the realm of the antibiotic armamentarium (1). They belong to the subclass of β-lactam antibiotics and exhibit high potency against multidrug-resistant (MDR) pathogens that infect adults and neonates (2). Overuse/misuse of this antibiotic in clinical settings promoted the emergence of carbapenem-resistant “superbugs” which produce “carbapenem-hydrolysing enzymes or carbapenemases” (3). Carbapenemases are classified into two molecular classes depending on their structure and functional characteristics, such as serine carbapenemases and metallo-β-lactamases (MBLs) (4). Serine carbapenemases (Ambler class A & D enzymes), such as IMI, KPC, OXA-48, SME and NMC-A are characterised by the presence of a serine residue in their active site which facilitates enzymatic action; whereas MBLs (Ambler class B enzymes) viz IMP, VIM and NDM require zinc ions at the active site for catalytic activity and facilitate hydrolysis of wide range of β-lactam antibiotics (5). Of the different MBLs, New Delhi metallo-β-lactamase (*bla*_NDM_), a subclass B1 enzyme, has disseminated worldwide among different Gram-negative bacteria due to its promiscuous nature (6, 7).

Different clinically important carbapenemases, such as NDM, IMP, VIM, KPC and OXA-48-like carbapenemases exhibit varied activity against carbapenems, which is reflected through the range of minimum inhibitory concentration (MIC) values detected through routine susceptibility tests (8). NDM-1 catalyses the hydrolysis of a wide range of β-lactams antibiotics with high efficiency (9). Studies have indicated that a subset of Enterobacterales possessing different carbapenemases, such as IMP and KPC, exhibited low MIC (<4 mg/L) values for meropenem (10). This low-level resistance to carbapenems, particularly meropenem MICs frequently below clinical breakpoints, suggested inherent differences in the ability of KPC and OXA-48-producing isolates to cleave meropenem (11, 12). Studies on class A carbapenemases (KPC) suggest that activity against carbapenems imposes specific spatial requirements on the class A β-lactamase active site that are necessary to orient bound carbapenems for hydrolysis. This, in turn, allows these enzymes to avoid inhibitory interactions with carbapenems compared to cephalosporins (13). These differences impair the ability of certain carbapenemase-producing isolates to efficiently hydrolyse meropenem, resulting in lower MIC values that fall within potentially treatable ranges (10).

In contrast, most *bla*_NDM_*-*positive (*bla* ^+ve^) isolates exhibit MICs for carbapenems in moderate to high ranges (16-128 mg/L) (6, 14). Rarely, exceptions have been noted where, despite the production of NDM, MIC values are much lower than the expected range (15–18). In this study, we attempted to understand the reasons for the low MIC values observed in isolates that possess *bla*_NDM_. The treatment of infections caused by NDM-producing bacteria in patients is primarily dependent on the results of *in vitro* antimicrobial susceptibility tests of pathogens against different antibiotics (19). Several studies have shown that, in rare cases, the clinical predictive values of these *in vitro* tests may not be correct (19). There may be several reasons for this mismatch, both dependent on the pathogen and the host (19). Given this, the study also examined whether isolates with *bla*_NDM_ and low MIC values for carbapenems could be effectively treated with meropenem. This was assessed through *in vitro* time-dependent bactericidal assays and *in vivo* experiments using a neutropenic mouse infection model.

## Materials and methods

### Bacterial isolates and antibiotic susceptibility

A total of 321 isolates were screened by PCR for the presence of *bla*_NDM_ collected between 2008-2019 from the blood of septicaemic neonates admitted in a neonatal intensive care unit of a tertiary care hospital in Kolkata. Among them, 123 (38.3%) were *bla* ^+ve^, which includes 29 *E. coli* (36.7%, 29/79) and 94 *K. pneumoniae* (38.8%, 94/242). Most of these isolates displayed meropenem MIC values in the range of 4-64 mg/L. In contrast, a few *bla* ^+ve^ isolates: one *E. coli* (EN5076) and five *K. pneumoniae* (EN5136, EN5137, EN5139, EN5144, EN5396) had low MIC values for meropenem (≤2 mg/L) based on current CLSI breakpoint interpretive guidelines (20).

Antibiotic susceptibility was assessed by the conventional disk diffusion assay (21). Antibiotic susceptibility for different antibiotics, such as amikacin, aztreonam, cefotaxime, cefoxitin, ciprofloxacin, colistin, gentamicin, meropenem, piperacillin, tigecycline, and trimethoprim-sulfamethoxazole (BD Diagnostics, Franklin Lakes, NJ, USA) was assessed by conventional disk diffusion assay. MIC values of carbapenems, i.e., meropenem, ertapenem, imipenem, and doripenem (Sigma-Aldrich, Germany), were evaluated by broth microdilution (20, 22). PCR-based screening of resistance determinants was carried out (Supplementary text). PCR-based screening of different resistance determinants such as, carbapenemases (*bla*_NDM_, *bla*_KPC_, *bla*_OXA-48 like,_ *bla*_IMP_, *bla*_VIM_)(23), β-lactamases (*bla*_SHV_, *bla*_TEM_, *bla*_OXA-1_, *bla*_CTX-M_)(24, 25), AmpCs (*bla*_MOX_, *bla*_CMY_, *bla*_DHA_, *bla*_ACC_, *bla*_MIR/ACT_, *bla*_FOX_) (26), 16S rRNA methylase-encoding genes (*armA, rmtA*, *rmtB*, *rmtC*, and *rmtD*) (27), and plasmid-mediated quinolone resistance genes (*qnrA*, *qnrB*, *qnrS*, *qnrC*, *qnrD*, *aac(6’)-Ib-cr*, *qepA*, *oqxA*, *oqxB*) (28) were carried out.

### Transmissibility of *bla*_NDM_

Conjugation for *bla*_NDM_*-*harbouring plasmids was done with *E. coli* J53 Az^r^ strain as recipient, and transconjugants (TCs) were screened on plates supplemented with 8 mg/L cefotaxime and 100 mg/L sodium azide (Sigma-Aldrich, St. Louis, USA). Electro-transformation was carried out for isolates in which conjugation was unsuccessful. *bla* ^+^ transformants (TFs) were selected on LB agar (BD Diagnostics, Franklin Lakes, NJ, USA) supplemented with ertapenem (0.5mg/L).

The presence of *bla*_NDM_ was confirmed, and other resistance determinants were detected in transconjugants (TCs) by PCR (29). Plasmid replicon types were identified in wild-type isolates along with their TCs or TFs using PCR-based replicon typing (PBRT) (Diatheva, Italy) (30).

### Sanger sequencing of *bla*_NDM_ and whole**-**genome sequencing (WGS)

*bla*_NDM_ amplicons (825 bp) were sequenced in an automated DNA sequencer (Applied Biosystems, Perkin Elmer, USA)(31). Pulsed-field gel electrophoresis (PFGE) depicted that four *K. pneumoniae* isolates were clonal [Figure S1]. All distinct isolates (EN5076 and EN5396) and two representative isolates (EN5137 and EN5139) from the clonal cluster were processed for WGS.

Library preparation for short-read sequencing was executed by Ion Xpress Plus fragment library kit (Thermo Fisher Scientific, Massachusetts, USA), and genomes were sequenced on Ion 540 chip with the Ion S5 system (Thermo Fisher Scientific, Massachusetts, USA). High-quality raw reads were assembled into contigs using SPAdes (v3.13.1) with default parameters (32, 33). Using the short-read data, different downstream analyses were carried out (Supplementary text) (34).

### *bla*_NDM_ expression by quantitative real-time PCR (qRT-PCR)

The mRNA expression of *bla*_NDM_ was studied for all isolates using a qRT-PCR assay. Cellular mRNA was extracted from 1 ml (∼10^8^ cells/ml) bacterial cultures, supplemented with cefotaxime (8 mg/L). Two micrograms of RNA was reverse transcribed into cDNA using RevertAid RT Reverse Transcription Kit (Thermo Fisher Scientific, Massachusetts, USA) according to the manufacturer’s instructions. Quantification of *bla*_NDM_ expression was performed with *bla*_NDM_ primer pair (NDM-F 5’-GGGCAGTCGCTTCCAACGGT-3’ and NDM-R 5’-CGACCGGCAGGTTGATCTCC-3’) and SYBR^TM^ Green PCR Master Mix in StepOne Plus Real-Time PCR System (Applied Biosystems, USA) (35). Expression of *bla*_NDM_ was normalised against the housekeeping gene, *rpsL* (small ribosomal subunit protein S12) to ensure consistency across the different samples (35, 36). Since all study isolates exhibited lower MIC values for meropenem (0.05-2 mg/L), another *bla* ^+ve^ *E. coli* isolate (EN5374) with a high MIC value for meropenem (64 mg/L) was considered as an endogenous control (expression = 1). Relative mRNA expression levels were determined using 2 ^-ΔΔCt^ method (Supplementary text).

### Mutation analysis of *bla*_NDM_ promoter region

The upstream of *bla*_NDM_ is consistently associated with an insertion sequence (IS) element known as IS*Aba125*, whose right inverted repeat (IRR) acts as the functional promoter of *bla*_NDM_ and controls the gene expression. This promoter features –35 box (TTGAAT), –10 box (TACAGT) for RNA polymerase binding sites and two putative transcription regulators, such as ArcA (TCATGTTT) and ArgR2 (CATATTTT) derived from the start codon of *bla*_NDM-1_(37, 38). The genetic environment surrounding *bla*_NDM_ (up to 126 base pairs upstream) was examined for all isolates (EN5076, EN5137, EN5139, and EN5396) using WGS data. The analysis was performed with SnapGene Viewer and compared to the *bla*_NDM-1_ nucleotide sequence (Accession FN396876) available in GenBank.

### Quantification of absolute copy numbers of *bla*_NDM_

Absolute quantification of *bla*_NDM_ gene copies was performed using droplet digital PCR (ddPCR) in study isolates exhibiting low meropenem MICs (0.5–2 mg/L), and the results were compared with two clinical *bla* ^+ve^ isolates showing elevated meropenem MIC values (16 and 64 mg/L). The ddPCR assay was carried out using the same primer pairs (*bla*_NDM_ and *rpsL*) used in the *bla*_NDM_ expression study (35, 36). The absolute count of *rpsL* copies was also determined to ensure consistency across the isolates.

Genomic DNA was isolated from EN5076, EN5144, and EN5396 (meropenem MIC: 0.5-2 mg/L); EN5374 and EN5180 (meropenem MIC: 64 and 16 mg/L) using the Wizard Genomic DNA Purification Kit (Promega, Madison, US) following the manufacturer’s instructions. All reagents, instruments and software used in this assay were purchased from Bio-Rad (Bio-Rad Laboratories, Hercules, CA, USA). Primer binding efficiencies for both primer pairs were calculated (Figure S2). A specific range of C_T_ values (∼24-26), which corresponded to 10 ng/µL of genomic DNA, was used as template DNA in ddPCR to ensure that both the positive and negative droplets and the counts come within the dynamic range of the instrument. For ddPCR, 20µL of reaction mixtures containing 2X EvaGreen ddPCR supermix, primers (0.25 µM) were included. Samples were distributed in a 96-well plate and loaded into the machine. Afterwards, automated droplet generation oil for EvaGreen and samples were loaded in DG32 automated droplet generator cartridge for the generation of droplets. The entire droplet emulsion volume (40μL) was further loaded into another 96-well PCR plate, heat-sealed with pierceable foil (PX1 PCR Plate Sealer) and placed in a thermal cycler following PCR conditions at 95°C-5 min, 39 cycles of denaturation at 95°C-30 s and elongation at 60°C-1 min, followed by signal stabilization at 4°C-5 min and 90°C-5 min and then cool down at 4°C. For all steps, a ramp rate of 2°C/s was used to ensure that each droplet reached the correct temperature. The PCR-generated droplets were read with the QX200 droplet reader and analysed with Quanta Soft droplet reader software, version 1.6.6.0320.

### *In vitro* time**-**dependent bactericidal activity of meropenem

Time-kill assay was performed to assess the bactericidal effect of meropenem on all study isolates for 72-hours. Meropenem concentration (8 mg/L) used in this experiment reflected the mean steady-state concentrations of nonprotein-bound drugs in humans (39). A single bacterial colony was inoculated into 3 mL Mueller-Hinton broth (BD Diagnostics, Franklin Lakes, NJ, USA) and incubated overnight at 37°C with shaking at 150 rpm. The next day, 50 µL of overnight culture was transferred to 5 mL fresh MHB and further incubated at 37°C to obtain a starting inoculum of approximately 5×10^5^ cfu/mL. Aliquots of bacterial cultures were collected at 0, 1, 2, 4, 6, 24, 48 and 72-hours, spread on LB agar, incubated at 37°C, followed by counting of bacterial colonies. The bactericidal effect of meropenem was defined as a ≥3 log_10_ decrease in cfu/mL after 24 hours compared to the initial inoculum (40).

### In vivo study

Female BALB/c mice were divided into three groups, with four mice in each group. Group 1 served as 0-hour control (infected mice sacrificed after 2-hours), group 2 served as a treatment group (infected and treatment initiated after 2-hours, sacrificed after 24-hours) and group 3 served as a 24-hours untreated group (infected mice sacrificed after 24-hours). Neutropenia (neutrophil count ≤100/mm3) was induced by intraperitoneal injection of cyclophosphamide (MP Biomedicals, California) 5 days before bacterial infection (41). Each posterior thigh muscle of the mice was challenged by an intramuscular injection of 100 µl of exponentially growing bacterial suspension (10^7^ cfu/mL) of EN5076 (*E. coli*) and EN5139 (*K. pneumoniae*).

In group 2, meropenem treatment was administered 2 hours after the bacterial infection at a dose of 100 mg/kg, given intraperitoneally, which simulated the human exposure [1 g of meropenem at every 8 hours (1 g q8 h 3-h)] (41). Group 1 mice were harvested at the beginning of dosing to serve as 0-hour untreated controls (∼baseline bacterial count). Group 2 and group 3 mice were sacrificed after 24-hours. Bacterial burden was quantified as colony-forming units (cfu) determined from posterior thigh homogenates and vital organs (lungs, liver and kidneys). The *in vivo* efficacy of meropenem was assessed by comparing the reduction in bacterial density (log_10_ cfu) in the thigh muscle of group 2 mice (meropenem-treated mice) from the initial bacterial load measured in group 1 mice (0 hours control group) [Supplementary text, Figure S3].

## Results

### Characteristics of the isolates

Six isolates were selected based on the presence of *bla*_NDM_ gene and MIC values in the range of 0.5-2 mg/L for meropenem. The isolates were multidrug-resistant (MDR), conferring resistance towards different antibiotics such as amikacin, aztreonam, cefotaxime, cefoxitin, ciprofloxacin, gentamicin, piperacillin, trimethoprim-sulfamethoxazole, and susceptible to meropenem, colistin and tigecycline (Table 1). Four isolates (EN5136, EN5137, EN5139 and EN5144) were clonal based on PFGE and had MIC values of 2 mg/L for meropenem (Figure S1). However, other isolates (EN5076 and EN5396) showed meropenem MIC values of 0.5 mg/L (Table 1).

**Table 1.**
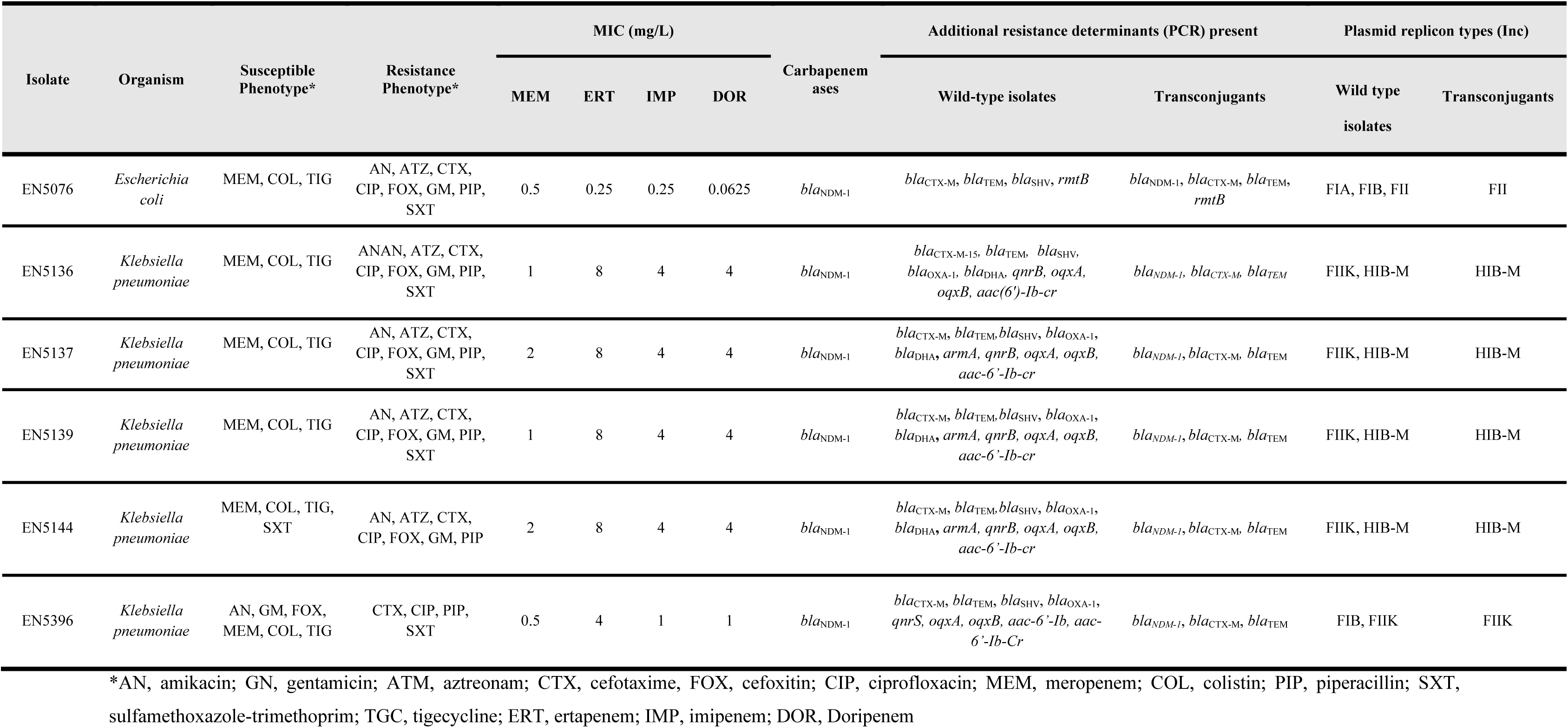
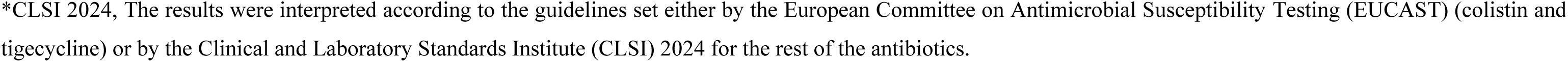
Antibiotic susceptibility, minimum inhibitory concentrations (MICs) of different carbapenems, with prevalence of resistance determinants among the wild-type isolates and transconjugants.

| Isolate | Organism | Susceptible Phenotype* | Resistance Phenotype* | MIC (mg/L) |  |  |  | Carbapenem ases | Additional resistance determinants (PCR) present |  | Plasmid replicon types (Inc) |  |
| --- | --- | --- | --- | --- | --- | --- | --- | --- | --- | --- | --- | --- |
|  |  |  |  | MEM | ERT | IMP | DOR |  | Wild-type isolates | Transconjugants | Wild type isolates | Transconjugants |
| EN5076 | *Escherichia coli* | MEM, COL, TIG | AN, ATZ, CTX, CIP, FOX, GM, PIP, SXT | 0.5 | 0.25 | 0.25 | 0.0625 | *bla*<sub>NDM-1</sub> | *bla*<sub>CTX-M</sub>, *bla*<sub>TEM</sub>, *bla*<sub>SHV</sub>, *rmtB* | *bla*<sub>NDM-1</sub>, *bla*<sub>CTX-M</sub>, *bla*<sub>TEM</sub>, *rmtB* | FIA, FIB, FII | FII |
| EN5136 | *Klebsiella pneumoniae* | MEM, COL, TIG | ANAN, ATZ, CTX, CIP, FOX, GM, PIP, SXT | 1 | 8 | 4 | 4 | *bla*<sub>NDM-1</sub> | *bla*<sub>CTX-M-15</sub>, *bla*<sub>TEM</sub>, *bla*<sub>SHV</sub>, *bla*<sub>OXA-1</sub>, *bla*<sub>DHA</sub>, *qnrB*, *oqx**A*, *oqx**B*, *aac*(6')-Ib-cr | *bla*<sub>NDM-1</sub>, *bla*<sub>CTX-M</sub>, *bla*<sub>TEM</sub> | FIIK, HIB-M | HIB-M |
| EN5137 | *Klebsiella pneumoniae* | MEM, COL, TIG | AN, ATZ, CTX, CIP, FOX, GM, PIP, SXT | 2 | 8 | 4 | 4 | *bla*<sub>NDM-1</sub> | *bla*<sub>CTX-M</sub>, *bla*<sub>TEM</sub>, *bla*<sub>SHV</sub>, *bla*<sub>OXA-1</sub>, *bla*<sub>DHA</sub>, *armA*, *qnrB*, *oqx**A*, *oqx**B*, *aac*-6'-Ib-cr | *bla*<sub>NDM-1</sub>, *bla*<sub>CTX-M</sub>, *bla*<sub>TEM</sub> | FIIK, HIB-M | HIB-M |
| EN5139 | *Klebsiella pneumoniae* | MEM, COL, TIG | AN, ATZ, CTX, CIP, FOX, GM, PIP, SXT | 1 | 8 | 4 | 4 | *bla*<sub>NDM-1</sub> | *bla*<sub>CTX-M</sub>, *bla*<sub>TEM</sub>, *bla*<sub>SHV</sub>, *bla*<sub>OXA-1</sub>, *bla*<sub>DHA</sub>, *armA*, *qnrB*, *oqx**A*, *oqx**B*, *aac*-6'-Ib-cr | *bla*<sub>NDM-1</sub>, *bla*<sub>CTX-M</sub>, *bla*<sub>TEM</sub> | FIIK, HIB-M | HIB-M |
| EN5144 | *Klebsiella pneumoniae* | MEM, COL, TIG, SXT | AN, ATZ, CTX, CIP, FOX, GM, PIP | 2 | 8 | 4 | 4 | *bla*<sub>NDM-1</sub> | *bla*<sub>CTX-M</sub>, *bla*<sub>TEM</sub>, *bla*<sub>SHV</sub>, *bla*<sub>OXA-1</sub>, *bla*<sub>DHA</sub>, *armA*, *qnrB*, *oqx**A*, *oqx**B*, *aac*-6'-Ib-cr | *bla*<sub>NDM-1</sub>, *bla*<sub>CTX-M</sub>, *bla*<sub>TEM</sub> | FIIK, HIB-M | HIB-M |
| EN5396 | *Klebsiella pneumoniae* | AN, GM, FOX, MEM, COL, TIG | CTX, CIP, PIP, SXT | 0.5 | 4 | 1 | 1 | *bla*<sub>NDM-1</sub> | *bla*<sub>CTX-M</sub>, *bla*<sub>TEM</sub>, *bla*<sub>SHV</sub>, *bla*<sub>OXA-1</sub>, *qnrS*, *oqx**A*, *oqx**B*, *aac*-6'-Ib, *aac*-6'-Ib-Cr | *bla*<sub>NDM-1</sub>, *bla*<sub>CTX-M</sub>, *bla*<sub>TEM</sub> | FIB, FIIK | FIIK |

\*AN, amikacin; GN, gentamicin; ATM, aztreonam; CTX, cefotaxime; FOX, cefoxitin; CIP, ciprofloxacin; MEM, meropenem; COL, colistin; PIP, piperacillin; SXT, sulfamethoxazole-trimethoprim; TGC, tigecycline; ERT, ertapenem; IMP, imipenem; DOR, Doripenem
\*CLSI 2024, The results were interpreted according to the guidelines set either by the European Committee on Antimicrobial Susceptibility Testing (EUCAST) (colistin and tigecycline) or by the Clinical and Laboratory Standards Institute (CLSI) 2024 for the rest of the antibiotics.

Genome-based characterisation showed that *E. coli* isolate (EN5076) belonged to epidemic clone ST410 with serotype O78-H18. All *K. pneumoniae* isolates had the same serotype, K62; however, clonal isolates belonged to ST48, while the remaining isolate was ST101 (Table 2). All isolates possessed *bla*_NDM-1_ detected by Sanger Sequencing and WGS. Apart from *bla*_NDM-1_, isolates carried multiple resistance determinants that conferred resistance to β-lactams (*bla*_NDM-1_*, bla*_CTX-M-15_*, bla*_TEM-1B_*, bla*_SHV-1_*, bla*_OXA-1_*, bla*_DHA-1_), and also other antibiotics such as aminoglycosides [*armA, rmtB*, *rmtC, aac(6’)-Ib-cr, aac(3)-Iid*)], sulfonamides (*sul1*), trimethoprim (*dfrA12*), phenicols (*catA1, catB3*), fluoroquinolones (*qnrB1, qnrS1, oqxA, oqxB*), and efflux pumps [*qacE*, *msr(E)*, *mph(E), mph(A)*] detected through PCR and WGS (Table 1, Table 2). Mutations in chromosomal genes encoding DNA gyrase (*gyrA*) and Topoisomerase IV (subunit A-*parC* and subunit B-*parE*) were also found that conferred elevated fluoroquinolone resistance (Table 2).

**Table 2:**
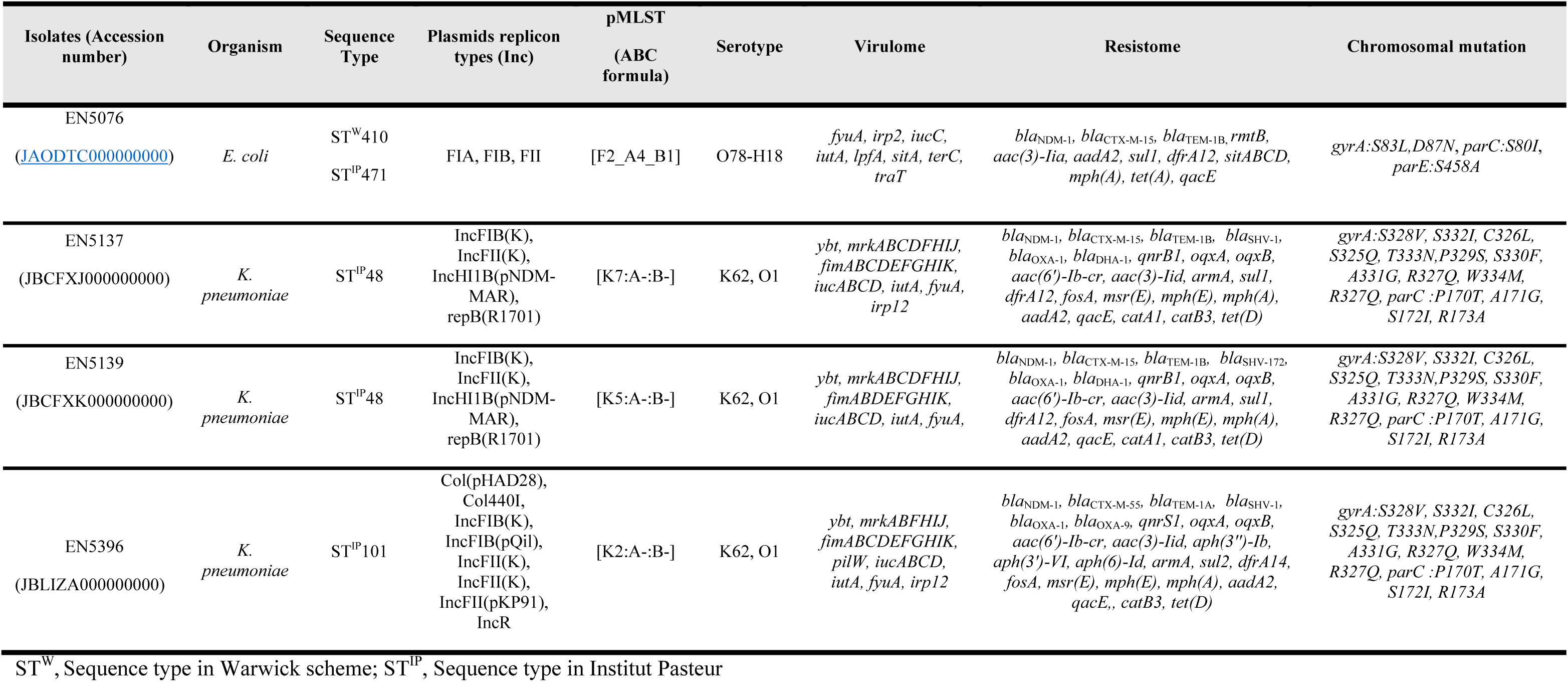
Genotypic characterisation of *bla*_NDM-1_-possessing *E. coli* and *K. pneumoniae* isolates based on whole-genome sequencing.

### Transmissibility of *bla*_NDM_

*bla*_NDM_ was successfully transferred via conjugation to the recipient *E. coli* J53 Az^r^ strain in four isolates (EN5136, EN5137, EN5139, EN5144) and to *E. coli* DH10B via transformation in two isolates (EN5076, EN5396), which confirmed its presence on plasmids. The plasmid replicon types associated with the *bla*_NDM_ include IncFII (EN5076), IncHIB-M (EN5136, EN5137, EN5139, EN5144) and FIIK (EN5396) (Table 1). Additional resistance determinants, such as *bla*_CTX-M_, *bla*_TEM_, *bla*_CMY_, *rmtB*, and *aac-(6’)-Ib-cr* were also found to co-transfer with *bla*_NDM_ (Table 1).

### Expression study of *bla*_NDM_

qRT-PCR assay showed downregulation in *bla*_NDM_ gene expression of study isolates (Figure 1). The C_T_ values for *bla*_NDM_ ranged from 21.9 to 25.6, indicating considerable differences in *bla*_NDM_ expression (Figure 1). Isolate EN5374 exhibited the highest *bla*_NDM_ expression (C_T_ = 20.9), serving as the endogenous control (meropenem MIC: 64 mg/L). Study isolates displayed *bla*_NDM_ fold change in a range of 0.27±0.02 to 0.48±0.12 compared to EN5374 (expression 1). Isolate EN5396 (meropenem MIC: 0.5 mg/L) demonstrated the lowest *bla*_NDM_ expression (C_T_ = 25.6) with a fold change of 0.064 compared to EN5374, i.e., ∼15-fold down-regulation. These findings suggest that *bla*_NDM_ gene expression can vary considerably among the isolates.

**Figure 1.**
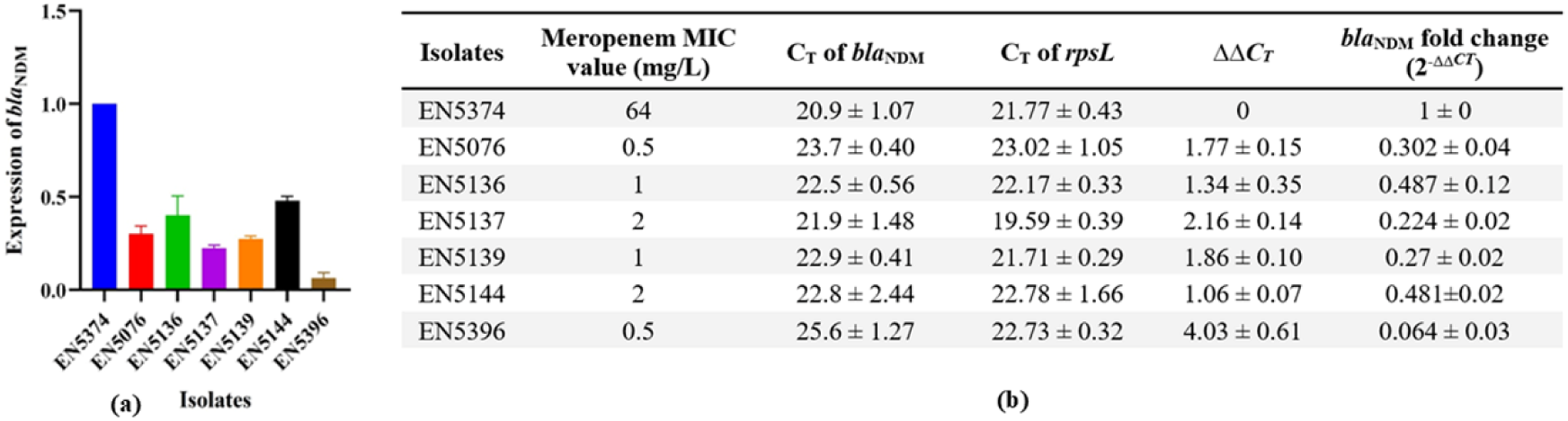
Graphical representation of *bla*_NDM_ expression in different study isolates relative to the reference isolate, EN5374 determined by real-time PCR (a) and relative expression of *bla*_NDM_ in different isolates (b). *EN5374 served as the endogenous control (expression = 1), exhibiting meropenem MIC value 64 mg/L. C_T_, cycle threshold; analysed by the 2^−ΔΔCT^ method. Data are presented as mean ± standard deviation (b).

### Analysis of the promoter region and genetic environment of *bla*_NDM_-_1_

The promoter region of *bla*_NDM_ was analysed from WGS. Alignment of the nucleotide sequences for all isolates revealed no mutations in the -35 box, -10 box, ArcA, and ArgR2 regions (Figure 2). Another upstream region, which serves as the ribosomal binding site/ RBS (-AAAGGAA) for translation, also showed no mutation. At the upstream, *bla*_NDM_ was always associated with IS*Aba125*, either full-length (3/4) or truncated (1/4) and *ble*_MBL_ was always found to be present downstream. Different downstream combinations were found in the individual isolates (Figure 2).

**Figure 2.**
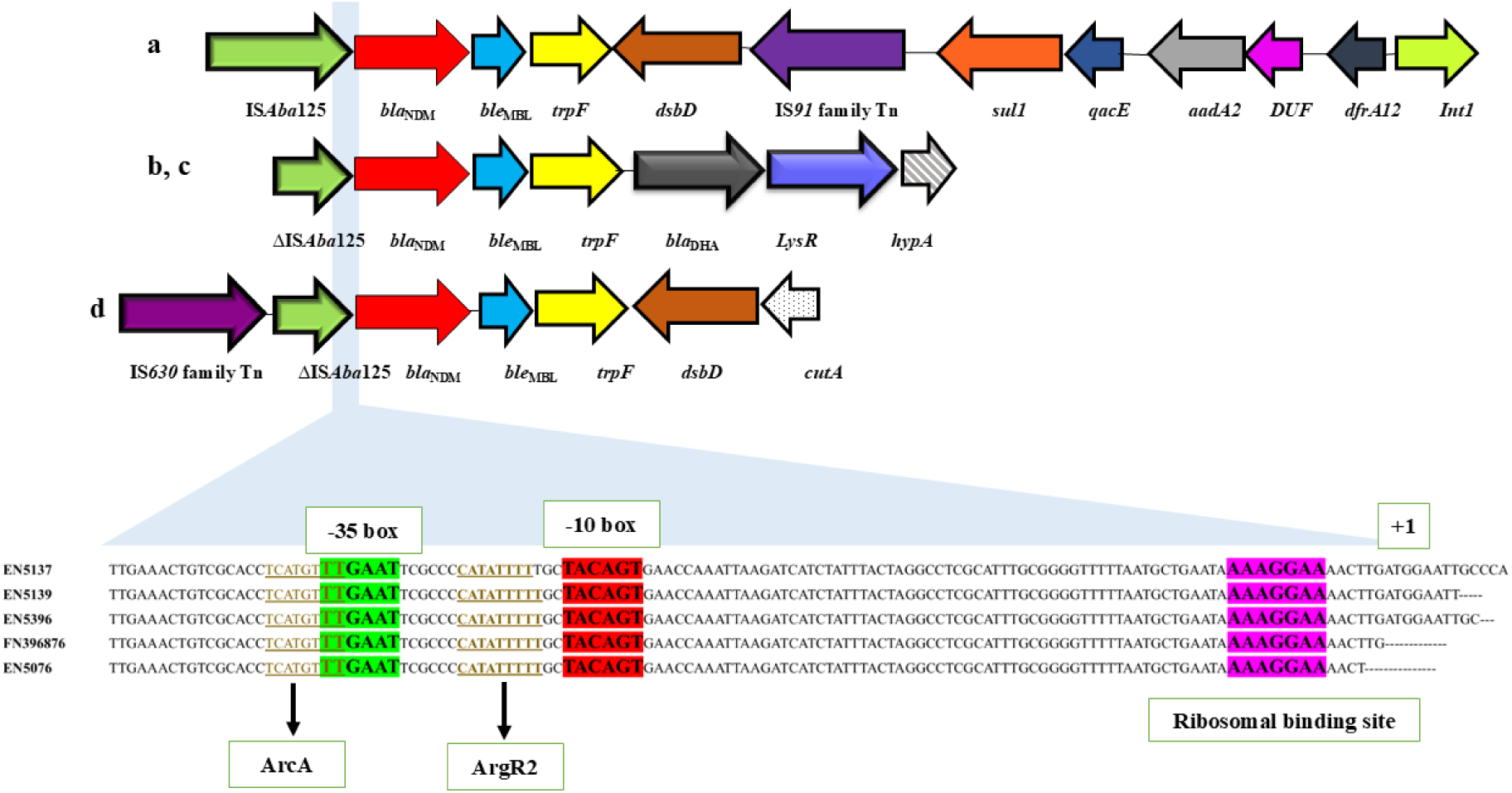
Schematic diagram of flanking regions of *bla*_NDM-1_ in study isolates and their promoter regions. The upstream region of *bla*_NDM_ in four study isolates, namely EN5076 (a), EN5137 (b), EN5139 (c), and EN5396 (d), was aligned with the *bla*_NDM-1_ sequence (Accession number FN396876) retrieved from NCBI for the detection of point mutations. The green highlighted region denotes the -35 box (-TTGAAT-), red highlighted region denotes the -10 box (-TACAGT-), and pink region denotes the ribosomal binding site of the promoter. The binding sites for the transcription regulators ArcA and ArgR2 are marked with brown underlining. The transcription start size is marked as +1 (ATG). *bla*_NDM-1_ gene sequence was downloaded from Genbank (https://www.ncbi.nlm.nih.gov/pathogens/refgene/#ndm-1).

### Copy number variation of *bla*_NDM_ among isolates

One-dimensional amplitude (1D) plot showed distinct clusters of positive and negative droplets of target genes by QuantaSoft droplet reader (Figure 3). Isolates having low MICs for meropenem, such as EN5076 (*bla*_NDM_ in IncFII) and EN5144 (*bla*_NDM_ in IncHIB-M), had a lower number of *bla*_NDM_ copies (6.3 × 10^4^ to 9.9 × 10^4^ copies/ul), suggesting low plasmid copies compared to isolates, EN5374 (7.1 × 10^5^ copies/uL; *bla*_NDM_ in IncFII) and EN5180 (8.2 × 10^5^ copies/uL; *bla*_NDM_ in IncHIB-M), having higher MIC values for meropenem (16-64 mg/L). Absolute quantification of *rpsL* revealed a range of approximately (6.3-9) × 10⁵ copies/µL across all samples. Therefore, the copy number of *bla*_NDM_ was correlated with reduced NDM expression detected through qRT-PCR, corroborated with lower meropenem MIC. Isolate EN5396 had an extremely low number of *bla*_NDM_ copies (45.11 copies/μL), and expression analysis was also 15-fold down-regulated (Figure 3), which indicated fewer gene copies and less enzyme production. Therefore, a reduction in *bla*_NDM_ gene dosage is directly proportional to the low meropenem MIC values of the isolates.

**Figure 3.**
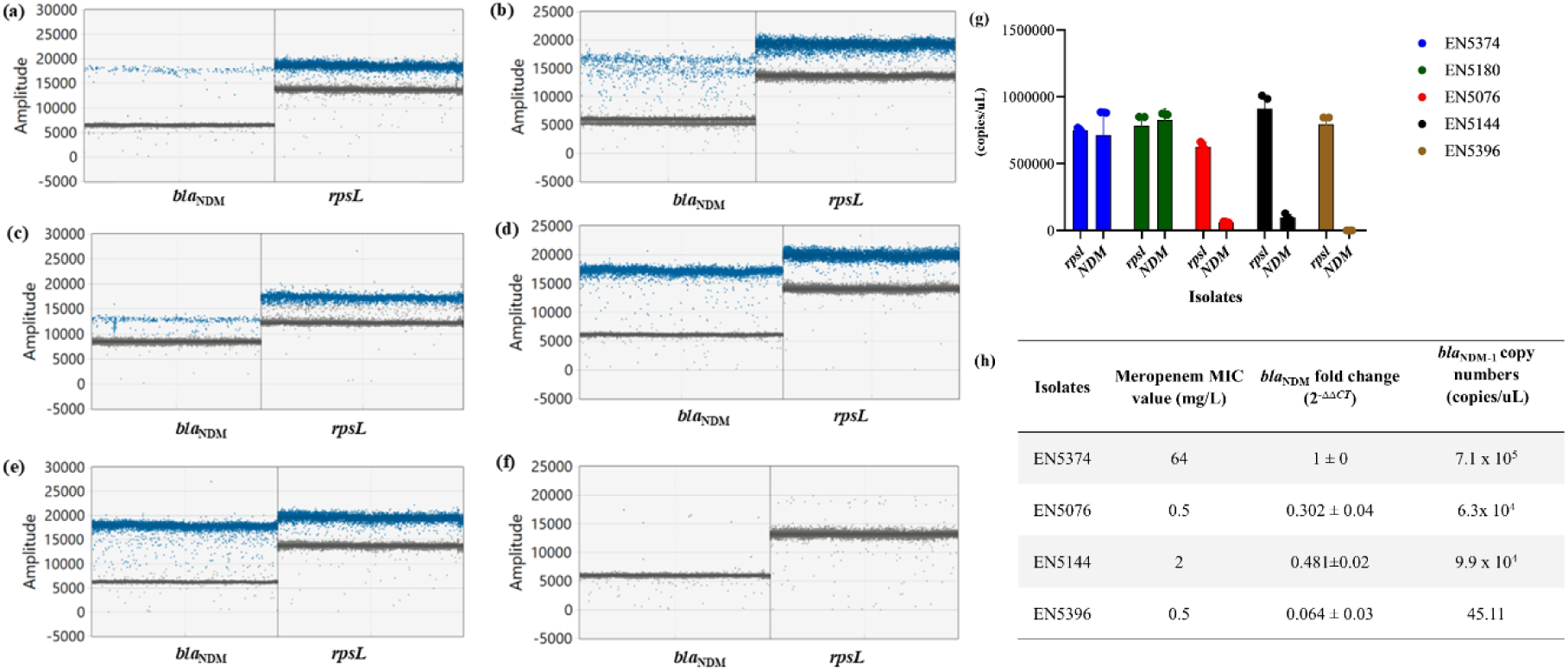
Absolute quantification of *bla*_NDM_ copies (copies/μL) in study isolates detected through droplet digital PCR (ddPCR). 1D-Amplitude plot showing droplets in blue dots (*bla*_NDM_^+ve^ & *rpsL*^+ve^) and droplets in grey dots (*bla* ^-ve^ & *rpsL*^-ve^) in separate panels for different study isolates [with meropenem MIC values 0.5-2 mg/L, EN5076 (a), EN5144 (b) and EN5396 (c)]; [meropenem MIC values ≥32 mg/L, EN5374 (d) and EN5180 (e)]; and non-template control (NTC), for detection of primer-dimer formation and DNA contamination (f). Graphical representation of *bla*_NDM_ copy numbers, with data expressed as mean ± standard deviation (g). Summary of study isolates with *bla*_NDM_ expression and absolute copy numbers (h). The housekeeping gene *rpsL* was used to normalise the data across different samples.

### *In vitro* time-dependent bactericidal assay

Bactericidal activity was observed for all isolates using meropenem as a single antibacterial agent at a concentration of 8 mg/L. Most isolates exhibited >3 log_10_ decrease in cfu/mL over 24-hour experimental period. EN5374 was used as a positive control due to its higher MIC value (64 mg/L) for meropenem. The rate of bacterial killing varied between isolates; isolate EN5137 showed a rapid decline in viable counts. Initial reductions of cfu (> 2 log_10_ cfu/mL), were noted during first two hours of all isolates in time-kill experiments. Marked regrowth was only observed in the case of isolate EN5374, which demonstrated higher resistance or tolerance to meropenem. No regrowth was noted for any of the tested isolates after 24 hours (Figure 4).

**Figure 4.**
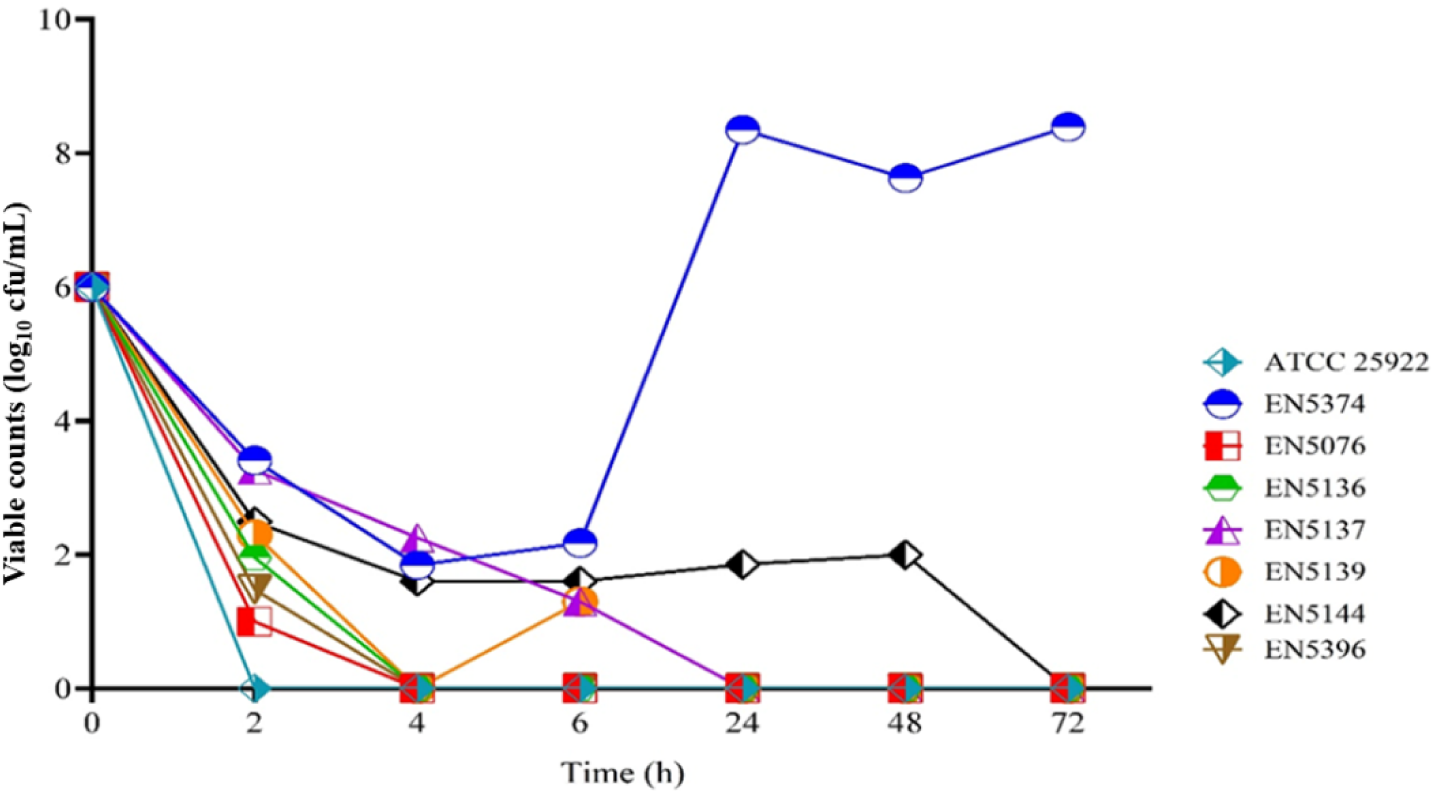
Time-kill curves showing *in vitro* bactericidal activity of meropenem at a concentration of 8 mg/L against study isolates. The *E. coli* ATCC 25922, susceptible to all antibiotics, was used as a negative control; another clinical isolate, EN5374, had a meropenem MIC of 64 mg/L and served as a positive control.

### In vivo assay

At the start of dosing, both isolates EN5076 and EN5319 displayed a mean bacterial burden of 7.15 ± 0.65 and 7.81 ± 0.12 log_10_ cfu/mL per thigh, respectively, for group 1 mice (0-hour untreated control). After 24 hours, the bacterial burden increased to 11.22 ± 0.15 log_10_ cfu/mL for EN5076 and 10.59 ± 0.31 log_10_ cfu/mL for EN5139 in group 3 mice (24 h untreated control). In contrast, meropenem-treated mice (group 2) showed a mean bacterial burden of 3.72 ± 0.17 log_10_ cfu/mL for EN5076, with all meropenem-treated mice surviving to the 24 h sampling point. Meropenem treatment with a 1-g dose (100 mg/kg for mice) resulted in an approximate 4 log cfu reduction for isolate EN5076, which had a MIC value for meropenem of 0.5 mg/L. For EN5319, group 2 mice exhibited a mean bacterial burden of 7.55 ± 0.08 log_10_ cfu/mL. Although this represented a reduction compared to the 0-hour control, meropenem did not demonstrate efficacy in the thigh muscle for this particular isolate, which had meropenem MIC of 2 mg/L. When comparing the treatment effects of meropenem on organs such as the lungs, liver, and kidneys, a significant reduction in bacterial burden was observed in group 2 mice compared to group 3 for both isolates (Figure 5).

**Figure 5.**
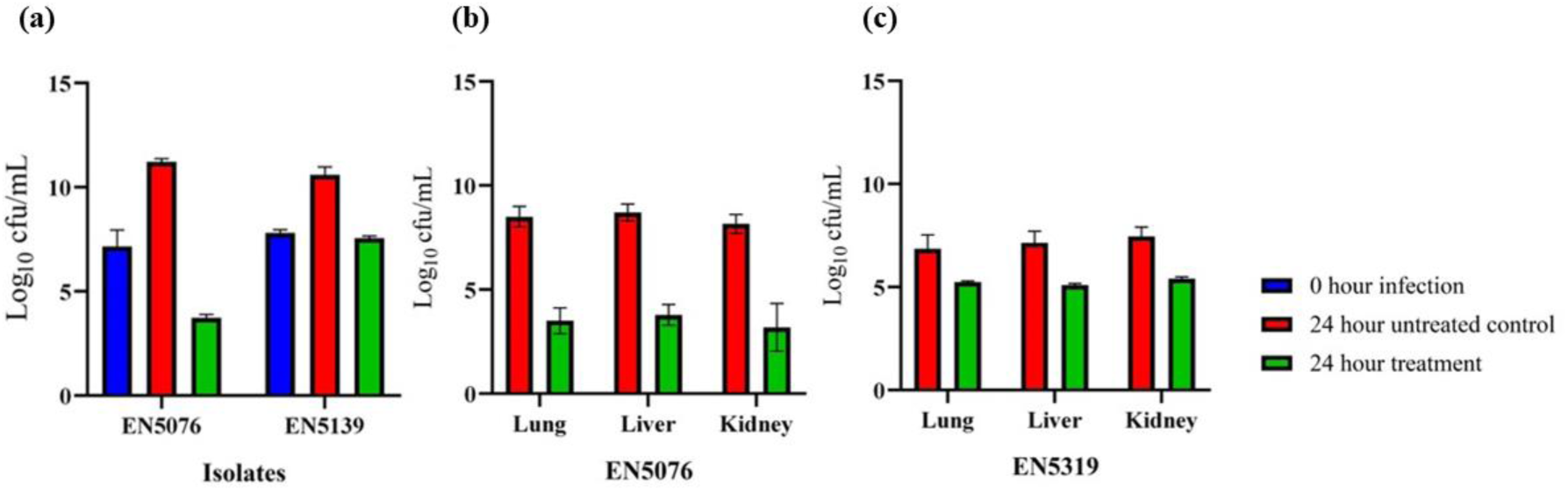
*In vivo* antibacterial activity of meropenem (100 mg/kg) in a neutropenic thigh infection model against *bla*_NDM-1_ positive clinical isolates EN5076 and EN5139. Bacterial load was measured in thighs (a), liver, kidney and lungs after 24 hours for both EN5076 (b) and EN5139 (c). Data are presented as mean ± standard deviation, based on values obtained from 3 mice in each group (group 1: 0-hour infection, group 2: 24-hour untreated control and group 3: 24-hour treatment).

## Discussion

NDM is an enzyme that can bind and hydrolyse a broad range of β-lactam antibiotics, including carbapenem, meropenem (6, 7). NDM-1 has high level of catalytic efficiency for varied substrates compared to other carbapenemases, which is normally reflected through moderate to elevated MIC values (42, 43). However, few studies have reported NDM-possessing isolates with low MIC values (15–18). This phenomenon had generally remained unexplored because it was infrequently observed. Our collection of 129 *bla* ^+ve^ *E. coli* and *K. pneumoniae* isolated during 2008-2019 showed MIC values in the range of 4-64 mg/L, which is in agreement with earlier studies (7, 8, 15). The exception to this particular observation was the presence of 6 isolates which had much lower MIC values in the range of 0.5-2 mg/L. These isolates carried multiple resistance determinants that conferred resistance to different antibiotics.

Initial analysis of the *bla*_NDM-1_ gene in six study isolates through both Sanger sequencing and WGS revealed an intact gene (813 bp). Antinori *et al*. reported a mutant variant of *bla*_KPC_ with a deletion of six nucleotides exhibited low meropenem MIC value (1mg/L)(44). However, this was not the case in this study. As a next logical step, the expression of the *bla*_NDM_ gene was assessed through real-time PCR which showed that isolates with low MIC values demonstrated down-regulation of *bla*_NDM_ gene [fold change 0.2-0.5] compared to an isolate with high MIC value (64 mg/L, expression=1). Studies have shown that a mutation in the promoter region can lead to reduced or elevated expression of a gene. Ching Hei Phoebe *et al*. reported that the rearrangement of the promoter region by insertion of a truncated *bla*_OXA-10_ gene reduced NDM-1 production in a clinical *Enterobacter* spp. (17). A study showed that *bla*_NDM_ gene silencing was caused by a natural mutation altering the sequence of the ribosomal binding site (AAAG**<u>G</u>**AA→AAAG**<u>A</u>**AA), located upstream of *bla*_NDM_ involved with reduced MIC of meropenem (MIC = 1 mg/L) in *bla*_NDM-5_^+ve^ isolates (45). Another study demonstrated that the functional activity of *bla*_NDM_ promoter remains within the 120 bp region upstream of *bla*_NDM_ (37). Based on this, the analysis of *bla*_NDM_ promoter region up to 126 base pairs, including the ribosomal binding site in study isolates was done, and no aberration was observed.

*bla*_NDM_ is primarily borne on plasmids and conjugation experiment for the study isolates showed that the gene could be transferred through plasmids such as IncFII, HIB-M, FIIK.

Absolute copy number estimation of *bla*_NDM_ by ddPCR demonstrated that isolates with lower MIC values had a relatively lower copy number of *bla*_NDM_ (6.3 x 10^4^ -10 x 10^4^ copies/uL) compared to *bla*_NDM_-positive isolates with high meropenem MICs (7 x 10^5^ copies/uL) suggesting that the variation in mRNA expression across isolates could be attributed to differences in the number of the *bla*_NDM_ copies probably because of the difference in the copy number of the plasmids harbouring the gene (46, 47).

Meropenem is a last resort drug that has bactericidal activity against MDR Enterobacterales with a pharmacokinetic profile which is characterised by rapid absorption, high bioavailability, minimal metabolism, excellent tissue penetration, efficient renal excretion and a generally well-tolerated safety profile (48). Hence, the use of meropenem should remain under consideration in patients with serious infections and not ruled out only because of the presence of a carbapenemase. Earlier *in vivo* studies have shown the efficacy of meropenem against VIM-producing *E. coli* and imipenem against KPC-producing *K. pneumoniae* with MICs ≤ 2 mg/L (10). With these factors in mind, we also evaluated the therapeutic efficacy of meropenem against isolates carrying *bla*_NDM_ and exhibiting low MIC values. The time-dependent bactericidal activity of this antibiotic, both *in vitro* time-kill assay and *in vivo* mouse model showed promise. Time-kill assay demonstrated 95% bactericidal activity (≥3 log_10_ decrease in cfu/mL), and simulated human dosing with 1 g of meropenem in neutropenic mice infected with two study isolates also demonstrated bactericidal activity (log_10_ decrease in cfu/mL). For both the tested isolates, a significant reduction in cfu was noted in vital organs, such as lungs, liver and kidneys, but cfu reduction in thigh muscle was observed for one isolate, EN5076, but was not significant for the other isolate, EN5139. We hypothesised that in organs such as lungs, liver and kidneys, which are highly vascularised, the penetration/bioavailability of antibiotics is higher than that in thigh muscle, which is not highly vascularised (49). *In vivo* study showed that meropenem retained therapeutic efficacy against study isolates. However, since dosing interval was not assessed in the animal model that remains a limitation of the study.

In recent times, to decrease the time for diagnosis, the reliance on molecular methods such as real-time PCR, NGS platforms, and microarray systems, which rapidly detect specific antibiotic resistance genes, has been noted. In such cases where MIC values are not determined, the reliance is on the detection of genes such as *bla*_NDM_, *bla*_KPC_, *bla*_OXA-48_, etc. may lead to a decision where carbapenems are not used. A well-tolerated drug like meropenem will have reduced usage and other toxic alternatives (colistin, which is nephrotoxic) may be administered. The study shows that meropenem may be considered for the treatment of infections with isolates that have low MIC values despite the presence of a carbapenem-resistant gene. When therapeutic options are already limited, it is critical to decide whether a drug like meropenem can be administered to critically ill patients for isolates similar to study isolates.

## Acknowledgements

We would like to express our deepest thanks to the entire staff and clinicians at the Neonatology Department of SSKM and IPGME&R Hospital for providing the isolates. We gratefully acknowledge Dr. Hemanta Koley for his guidance in animal experiments. We also thank Dr. Wriddhiman Ghosh for his valuable suggestions to execute genome sequencing and assembly.

## Data availability statement

All of the genome data were submitted to the NCBI database under the BioProjects PRJNA548120 (https://dataview.ncbi.nlm.nih.gov/object/PRJNA548120) (EN5136, EN5137, EN5139, EN5144 and EN5396) and PRJNA801430 (https://www.ncbi.nlm.nih.gov/bioproject/PRJNA801430) (EN5076).

## Ethical approval

This study was carried out on archived isolates, so ethical clearance was not required. The Institutional Animal Ethics Committee (IAEC) reviewed and approved the study for animal experiments, proposal number PRO/187/-Nov2022-25.

## Funding

This study was supported by the Indian Council of Medical Research (ICMR) intramural fund.

A.B. was supported by the fellowship from the ICMR.

## Transparency declarations

None to declare.

